# Pylluminator: fast and scalable analysis of DNA methylation data in Python

**DOI:** 10.1101/2025.09.16.676547

**Authors:** Elio Fanchon, Benjamin Loire, Jean-Philippe Trani, Frédérique Magdinier, Anaïs Baudot

## Abstract

**Motivation:** Illumina Infinium BeadChip technology for DNA methylation analysis continues to expand, with the latest EPICv2 arrays targeting about a million loci. As data volumes from this technology continue to grow, there is increasing demand for more scalable data processing solutions. Meanwhile, Python has gained significant interest in bioinformatics for its efficiency, versatility, and widespread use in data science and machine learning. Yet, no comprehensive Python toolkit exists for Illumina methylation array analysis.

**Results:** We present Pylluminator, a Python implementation of essential analysis methods including pre-processing tools, quality control, differential methylation analysis, and visualizations. Based on the established R packages SeSAMe and ChAMP, Pylluminator provides a scalable, user-friendly toolkit for DNA methylation analysis.

**Availability and implementation:** Pylluminator is an open-source package under MIT license available at https://github.com/eliopato/pylluminator. It was developed using Python 3.12 and can be installed with pip. The documentation with thorough installation instructions and examples can be found at https://pylluminator.readthedocs.io

## Introduction

Epigenetics, a rapidly expanding field in biology, refers to the study of heritable changes able to modify the genome activity without alteration of the DNA sequence itself. Epigenetics involves chemical modifications of the DNA molecule, modifications of histone proteins around which DNA is wrapped to form chromatin, and the regulatory action of non-coding RNAs (Li, 2020).

DNA methylation, one of the key epigenetic chemical modifications of DNA, is a powerful genome regulation mechanism. It plays a causative role in the development of various pathologies, including cancer (Sharma et al., 2010), autoimmune disorders, neurological disorders or rare genetic diseases such as ICF (Immunodeficiency, centromeric instability and facial anomalies), or Tatton-Brown-Rahman syndromes among others (Moosavi and Ardekani, 2016). DNA methylation occurs when a methyl group is added to the DNA molecule on an adenine or cytosine base (Ratel et al., 2006). In mammals, DNA methylation mainly occurs on the cytosines of CpG dinucleotides and the majority of CpG dinucleotides (60% to 90%) are methylated. CpG-rich regions called ‘CpG islands’, which are frequently found in gene promoter regions, show variable methylation patterns (Bird, 1986). Methylation of CpG islands usually contribute to gene silencing. Methylation of repetitive DNA sequences also contribute to the maintenance of genome integrity.

Illumina Infinium Methylation BeadChips offer high-resolution tools to quantify genome-wide DNA methylation (Bibikova et al., 2005). Various bioinformatics packages, such as ChAMP (Tian et al., 2017), SeSAMe (Triche et al., 2013) or Minfi (Aryee et al., 2014), have been developed over the years to process and analyze Illumina Infinium BeadChips data. Unfortunately, these packages are being challenged by the increasing number of probes and samples in experiments. For instance, the latest Illumina Infinium BeadChips version for the human genome, EPICv2, targets over 930.000 loci, of which 99.5% are CpG sites.

With the exponential growth of interest in data science over the past decade, Python became one of the most used languages to handle large-scale datasets efficiently. An increasing number of bioinformatics tools are being developed or translated to Python, but there is no up-to-date tool available to fully process, analyze, and visualize Illumina methylation array data. We propose to fill the gap with Pylluminator, which implements the most common pre-processing methods, analysis algorithms, and data visualizations of DNA methylation (Fig. 1). Pylluminator is based on the reference R packages SeSAMe and ChAMP and provides a user-friendly, scalable, and reproducible toolkit for methylation data analysis, bridging the gap between bioinformatics and data science.

**Fig. 1.**
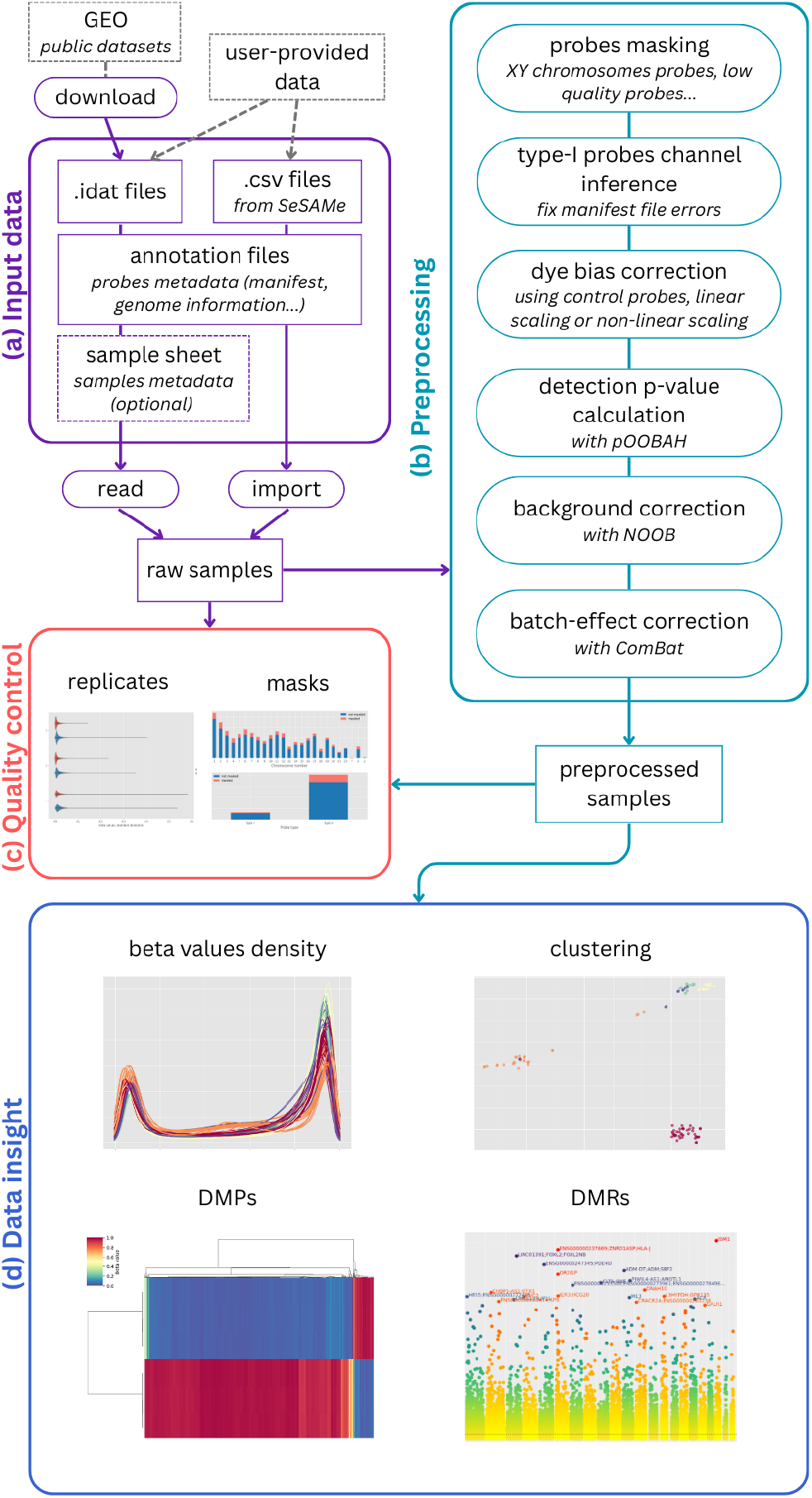
Flowchart of the main features of Pylluminator.

### Input data

Pylluminator handles all human (27k, 450k, MSA, EPIC, EPIC+, EPICv2), mammalian (Mammal40) and mouse (MM285) array versions, and supports different genome versions (hg38 and hg19 for human arrays, mm39 and mm10 for mouse arrays). The annotations corresponding to each array, which we have created from SeSAMe and Illumina annotation files, are automatically downloaded from https://github.com/eliopato/pylluminator-data when needed - but custom annotations can also be provided by the users. The annotation files include the manifest file that maps each probe’s Illumina ID to its metadata (e.g., color channel, location on the chromosome, probe type). The annotation files also include genome information files for data enrichment and visualizations (e.g., transcripts description, chromosome regions information).

As described in Fig. 1 (a), the input methylation data can be raw. *idat* files, as well as an existing SigDf object exported from SeSAMe as a .*csv* file. The raw .*idat* files can come from user-provided experimental data, or can be automatically downloaded from Gene Expression Omnibus (GEO) database by providing the samples GSM ids.

### Pre-processing

As recommended by SeSAMe, the pre-processing pipeline starts by masking unwanted probes (Fig. 1 (b)). Unwanted probes can be control probes, SNP probes, all non-CG probes, non-unique probes, probes that are known to have design issues (as defined by Zhou et al. (2016)), XY chromosome probes, as well as user-defined masks. The masking framework in Pylluminator makes it easy to exclude probes from the study, and masks are reversible at any time in the pipeline.

The second pre-processing step aims to infer the channel of type I probes. As detailed in the ‘Input Data’ section, probes metadata, including color channel for type I probes, are provided by the manifest files. Running the Type-I channel inference method before any further processing allows to correct potential errors in the annotation of type I probes.

Then, dye bias, a well-known issue in multichannel microarray experiments that is due to a difference in fluorescence intensity between the green (Cy3) and red (Cy5) channels, can be corrected with linear or nonlinear scaling.

Next, detection p-values are commonly used to assess the likelihood that a given probe signal corresponds to background fluorescence. We chose to integrate pOOBAH (P-value with Out-Of-Band Array Hybridization) (Zhou et al., 2018), a method that calculates detection p-values based on the empirical cumulative distribution function of overall Type-I out-of-band signal. Probes with detection p-values higher than a certain threshold are then masked (by default, 0.05). Zhou et al. (2018) also show that masking these probes can highly reduce technical variations, such as batch effect (Leek et al., 2010).

For overall background signal correction, Pylluminator implements NOOB, the background subtraction algorithm based on Normal-exponential deconvolution using Out-Of-Band probes, as defined by Ritchie et al. (2007) and Triche et al. (2013).

If a batch effect remains, Pylluminator also integrates the most widely used algorithm for correcting technical biases in microarray data, ComBat (Johnson et al., 2006). ComBat is based on an empirical Bayes framework, and handles low sample sizes and more than two batches at the same time.

### Samples quality control

Pylluminator includes comprehensive quality control (QC) tools to identify potential issues in methylation data. Users can evaluate key metrics such as probe detection rates by probe type (CG, CH, SNP), average signal intensity by probe design type (I or II) and channel (red or green), type I probes color channel balance, and beta values distribution by probe type. These tools can be used to assess the quality of raw input data and the effectiveness of the pre-processing pipeline.

To further diagnose technical variability, the replicate analysis plot helps detect batch effects by comparing technical replicates. Additionally, visualizing the distribution of masked probes per chromosome and by probe design type provides valuable insights into experiment quality (Fig. 1 (c)).

### Data insights

Beta-values are calculated for each probe using SeSAMe’s formula, defined as:

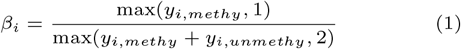

where *y*_*i,methy*_ and *y*_*i,unmethy*_ are the intensities measured by the *i*^*th*^ methylated and unmethylated probes, respectively.

To detect Differentially Methylated Probes (DMPs), we fit an Ordinary Least Squares (OLS) linear regression model to each probe’s beta values. Any metadata extracted from the sample sheet (e.g., age, sex, phenotype) can be used as predictor in the formula to describe the model. The resulting p-values and effect size are then used to identify the significant DMPs, after correcting for multiple testing using the Benjamini-Hochberg procedure (Benjamini and Hochberg, 1995). If the experimental design includes technical replicates, repeated measures, or any other source of random variability, a Linear Mixed Model (LMM) is used instead of OLS. In the LMM, predictors of direct interest (e.g., age, sex, phenotype) are modeled as fixed effects, while sources of random variability (e.g., patient ID, batch effects) are modeled as random effects. This accounts for within-group correlations and improves the accuracy of inference.

Differentially Methylated Regions (DMRs) are then identified by grouping neighboring probes with similar methylation patterns for a given predictor contrast (e.g., treated vs. control). Similarity is calculated based on the Euclidean distance between probes’ beta values. The distance cut-off used to separate the regions can be explicitly specified, or automatically set to the value at a specified quantile of all pairwise distances (by default, 0.5, i.e., the median). The significance of each DMR is then computed by combining the p-values of its constituent DMPs with Stouffer’s Z-score method (Stouffer et al., 1949).

The Copy Number Variation (CNV) calculation method implemented in Pylluminator relies on copy-number-normal samples to normalize probe signal intensity. By default, Pylluminator provides precomputed normal reference data for EPIC/hg38 and EPICv2/hg38 array versions, similar to SeSAMe’s implementation. For other array versions, users must provide their own reference data. Regardless of the array version, it is strongly recommended to use normal samples that closely match the biological and technical characteristics of the target samples (e.g., same tissue type, platform, and processing batch) to ensure accurate normalization.

For Copy Number Segmentation (CNS), Pylluminator employs the Circular Binary Segmentation (CBS) algorithm to identify genomic regions with homogeneous CNV. This approach efficiently detects breakpoints and segments the genome into intervals of consistent copy number variation.

## Results

We compared the major data processing and analysis functions of Pylluminator with their SeSAMe or ChAMP equivalents on a dataset of 24 EPIC samples from Laberthonniére et al. (2023). Computation time is evaluated by running the functions 10 times, on a laptop with 32 Gb RAM and an Intel Core i7 processor with 14 cores. Batch correction is applied using Sentrix ID as batch identifier, and samples’ phenotype as covariate. We compute the DMPs using the samples phenotype predictor (Control/FSHD1/FSDH2/BAMS). Results are presented in Table 1.

**Table 1.**
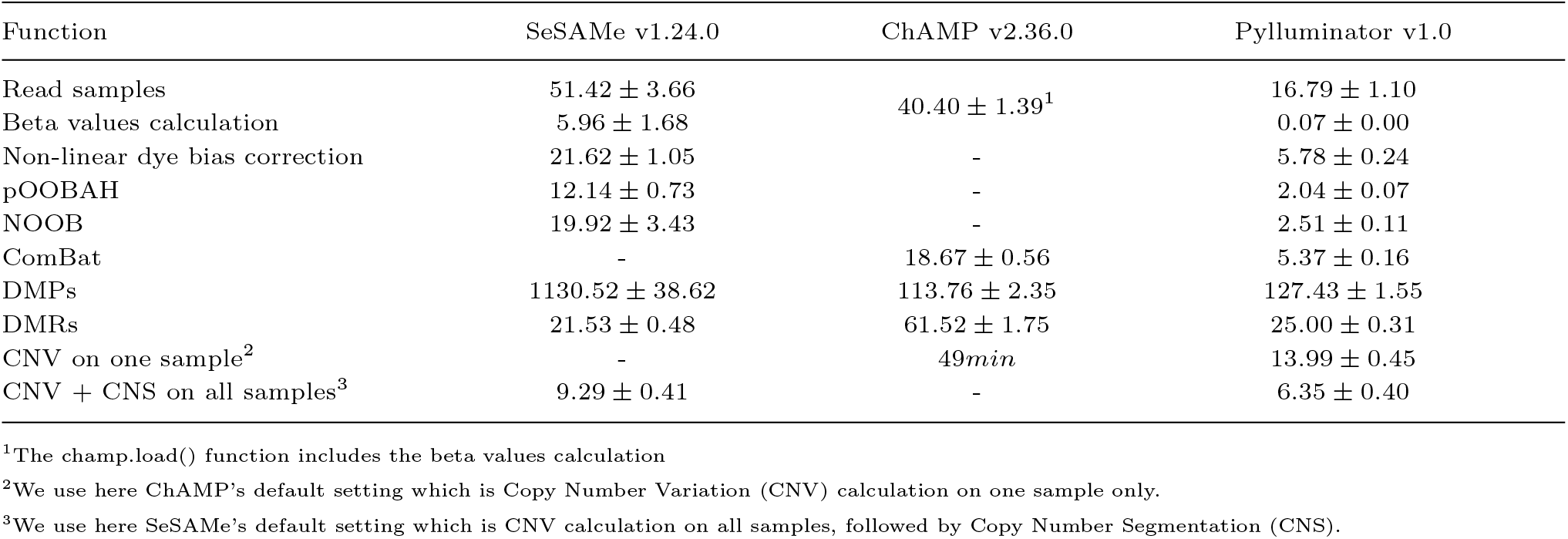
Comparison of average execution time *±* standard deviation (in seconds, unless otherwise stated) of the main functions.

## Code reliability

The code follows PEP8 standards and comes with extensive documentation, making it easy for external contributors to extend Pylluminator’s functionalities. The code and the data are hosted on GitHub at https://github.com/eliopato/pylluminator-data and https://github.com/eliopato/pylluminator.

To ensure consistency and reliability, 93% of the code is covered by unit tests that systematically check for regressions or unwanted changes, and compare the results of Pylluminator with the outputs of reference tools such as SeSAMe (Triche et al., 2013) and ChAMP (Tian et al., 2017).

## Acknowledgments

Pylluminator exists thanks to the growing Python bioinformatics community. We would like to thank the developers of the packages used in Pylluminator: methylprep for the .idat files parser, inMoose (Behdenna et al., 2023) for the implementation of the ComBat algorithm, and pyRanges (Stovner and Sætrom, 2020) for handling GenomicRanges.

We also thank the developers of the R packages used as reference, namely SeSAMe (Triche et al., 2013) and ChAMP (Tian et al., 2017).

Last but not least, we thank David Hirst and Youssef Boueimen for testing Pylluminator thoroughly and giving valuable feedbacks.

## Abbreviations

BAMS: Bosma Arhinia Microphthalmia Syndrome
CNS: Copy Number Segmentation
CNV: Copy Number Variation
DMP: Differentially Methylated Probe
DMR: Differentially Methylated Region
FSHD1 Type 1: FacioScapuloHumeral muscular Dystrophy
FSHD2 Type 2: FacioScapuloHumeral muscular Dystrophy
LMM: Linear Mixed model
NOOB: Normal-exponential Out-Of-Band
OLS: Ordinary Least Squares
pOOBAH: p-values Out-Of-Band Array Hybridization

